# Glandular quinoline-derivates protect crustacean woodlice from spider predation

**DOI:** 10.1101/2025.03.25.645209

**Authors:** Andreas Fischer, Regine Gries, Camila A. Roman-Torres, Anand Devireddy, Gerhard Gries

## Abstract

In evolutionary time, aquatic crustaceans colonized land and faced new terrestrial predators such as spiders and ants. Working with the crustacean woodlouse *Porcellio scaber*, we tested the hypothesis that the shift from aquatic to terrestrial habitat prompted the evolution of defensive metabolites against terrestrial predators. When attacked by a predator, *P. scaber* woodlice expel proteinaceous secretions from their tegumental glands. Analyses of gland secretion extracts by gas chromatography-mass spectrometry and by liquid-chromatography-tandem mass spectrometry revealed four metabolites: methyl 8-hydroxy-quinoline-2-carboxylate, methyl 8-hydroxy-4-methoxy-quinoline-2-carboxylate, methyl 8-(sulfooxy)quinoline-2-carboxylate, and methyl 4-methoxy-8-(sulfooxy)quinoline-2-carboxylate, the latter three being natural products not previously known. In behavioural experiments, *Steatoda grossa* spiders preyed on chemically undefended *Tenebrio molitor* beetles but avoided chemically defended *P. scaber*. When beetles were rendered chemically defended by topical applications of either *P. scaber* gland secretion extract or synthetic metabolites identified in these extracts, spiders rejected the beetles as prey. Our data support the hypothesis that the evolutionary transition of crustaceans from aquatic to terrestrial habitats prompted the evolution of defensive metabolites against terrestrial predators. We show that crustaceans, like many insects, are chemically defended against predators.

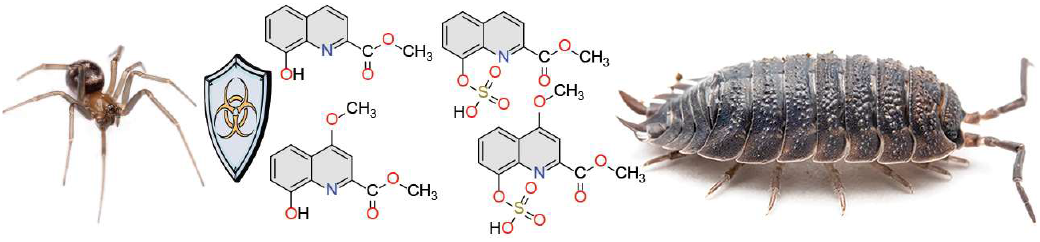

**Graphical abstract:** Under attack by the predatory spider *Steatoda grossa*, the crustacean *Porcellio scaber* releases four defensive metabolites: methyl 8-hydroxy-quinoline-2-carboxylate, methyl 8-hydroxy-4-methoxy-quinoline-2-carboxylate, methyl 8-(sulfooxy)quinoline-2-carboxylate, and methyl 4-methoxy-8-(sulfooxy)quinoline-2-carboxylate, the latter three previously unknown natural products.

## 1. Introduction

In the struggle for survival, predation has prompted the evolution of antipredator defences in prey [1]. Behavioural adaptations, such as fleeing or feigning death in response to a predator, are widespread across the animal kingdom [2]. Similarly, protective morphological traits, such as hard exoskeletons or spines, have evolved independently in various lineages throughout the tree of life [3,4]. Chemical defences by prospective prey entail the secretion of predator-repelling (foul-smelling) or predator-deterring (foul-tasting) metabolites [5].

Animals often employ a combination of anti-predator defences to enhance their survival [6]. Unlike behavioural and morphological defence traits, chemical defences remain largely unexplored [7,8] due, in part, to analytical obstacles for studying them [9]. Specifically, the vast diversity of behaviour-modifying metabolites, the difficulty of isolating them, and the complexity of identifying their molecular structure pose significant challenges [9]. Chemical defences evolved across many animal taxa, from protozoans to vertebrates, including all life-stages [5]. Defensive metabolites serve either in a pre-emptive or a reactive line of defence. Poisonous metabolites on salamander skin exemplify a pre-emptive chemical defence [10], whereas the foul-smelling secretions of a skunk threatened by a predator represent a reactive chemical defence [11]. The bioactive metabolites in these defences are diverse, ranging from complex enzymes in venoms [12] to simple formic acid in defensive sprays of ants [6]. Mostly insect-produced defence metabolites have been identified to date, while not a single defence metabolite produced by crustaceans has been identified [5,13]. This is astounding because crustaceans are close phylogenetic relatives of insects and are taxonomically diverse [14].

Crustaceans are mostly aquatic but polyphyletic “Oniscidea” isopods (woodlice) have colonized land [15,16]. In their adopted terrestrial habitats, early Oniscidea isopods faced new terrestrial predators, prompting the evolution of adaptive defence tactics [17]. These tactics entail behaviours such as grouping with reduced predation risk for individuals [18], scattering and fleeing, rolling into a ball (conglobating), or producing stridulatory sound when disturbed [17]. Besides behavioural adaptations, terrestrial isopods have evolved an array of morphological defences, including a hardened exoskeleton, spiny tergites, and morphs with aposematic coloration [17]. The key chemical defence of woodlice pertains to proteinaceous tegumental gland secretions [19] that coagulate within seconds of release and render the woodlice deterrent to potential predators [5,17,19,20] such as ants and spiders [20,21]. The common woodlouse, *Porcellio scaber* (Oniscidea: Porcellionidae), is particularly deterrent to predators [19,20].

We tested the hypothesis that the evolutionary transition of crustaceans from aquatic to terrestrial habitats prompted the evolution of defensive metabolites in tegumental gland secretions against terrestrial predators. We tested this hypothesis by identifying metabolites in tegumental gland secretions from *P. scaber* and by testing their behavioural effect on predatory spiders. As a generalist predatory spider, we used female false widows, *Steatoda grossa* (Araneae: Theridiidae), which may prey on *P. scaber* [22]. We predicted (1) that spiders prey more readily on chemically-undefended *Tenebrio molitor* beetles than on chemically-defended *P. scaber*, and (2) that spiders reject beetles rendered chemically-defended by treatment with synthetic *P. scaber* secretion metabolites.

## 2. Methods

### (a) Animal husbandry

*Porcellio scaber* woodlice were collected near Tsawwassen, British Columbia, Canada (49° 0’ 53.856’’ N, 123° 2’ 26.6532’’ W) and housed communally in a plastic bin (61 × 39.5 × 21 cm) fitted with mesh-covered holes for ventilation and filled partially with moist C-I-L® Black Earth Top Soil (Riviere-du-Loup, QC, Canada). Woodlice were provisioned with organic potatoes and fish food (Nutrafin basix Staple Food, Rolf C. Hagen Inc., Montreal, Canada) *ad libitum*. Adult *Tenebrio molitor* mealworm beetles were housed communally in a similar bin filled with wheat bran and provisioned with organic potatoes and fish food *ad libitum* [23]. Adult female *S. grossa* spiders were housed singly in translucent 300-mL plastic cups (Western Family, Canada) and fed four *Phormia regina* black blow flies per week [24]. All animals were kept at 22 °C under a reversed light cycle (12:12 h).

### (b) Collection of secretions from *P. scaber* tegumental glands

Defensive secretions from adult *P. scaber* were collected by immobilizing specimens with modelling clay (Craftsmart®, Michaels Stores Inc., Irving, TX, USA) and bringing a heated metal probe briefly into contact with the woodlouse’s tergites [19], thus stimulating the release of defensive secretions from their tegumental glands (suppl. video 1). Secretions (< 1 µL per woodlouse) were collected in 5-µL microcapillaries (Drummond Microcaps, Drummond Scientific Comp., Broomall, USA). The secretions from 83 *P. scaber* were pooled, dissolved in HPLC-grade methylene chloride (40 µL per secretion), and then concentrated under a gentle stream of nitrogen to achieve a final concentration of one secretion equivalent per one microlitre. Adult *P. scaber* replaced their compromised integument through moulting.

### (c) Chemical analyses of *P. scaber* tegumental glands secretions

In search for defensive metabolites, aliquots (2 µL) of secretion extract were analysed using a 5977A Mass Selective Detector (MSD) coupled to a 7890B gas chromatograph (GC) (Agilent, Santa Clara, USA) fitted with a DB-5 column (30 m × 0.25 mm ID, film thickness 0.25 µm). The injector port was set to 250 °C, the MS source to 230 °C, and the MS quadrupole to 150 °C. Helium was used as a carrier gas at a flow rate of 35 cm s^−1^, with the following temperature program: 40 °C held for 5 min, 10 °C min^−1^ to 280 °C (held for 20 min) [25].

In search for very polar metabolites that would not be detectable by GC-MS analyses [26,27], aliquots (2 µL) of secretion extract were further analysed using liquid chromatography with tandem mass spectrometry (LC/MS-MS). The system consisted of an Agilent 1200 LC fitted with a Spursil C_18_ column (30 mm × 3.0 mm, 3 µm; Dikma Technologies, Foothills Ranch, CA, USA) and a Bruker maXis Impact Ultra-High Resolution tandem TOF (UHR-Qq-TOF) mass spectrometer. Electrospray ionisation was set to positive (+ESI) at a gas temperature of 200 °C and a flow of 9 L/min. The nebuliser was set to 4 bar and the capillary voltage to 4200 V. The column was eluted with a 0.4 mL/min flow of a solvent gradient starting with 95% water and 5% acetonitrile, and ending with 100% acetonitrile after 4 min [28]. Identified metabolites were synthesized as outlined in the supplementary material.

### (d) Testing for defensive functions of *P. scaber* tegumental gland secretions and synthetic gland metabolites

Prediction 1 – spiders prey more readily on chemically-undefended *Tenebrio molitor* beetles than on chemically-defended *P. scaber* – was tested in experiments 1–4. In each experimental replicate a spider, food-deprived four weeks [29] to enhance her predatory responsiveness, was transferred to an inverted 300-mL cup and was offered as prey a *T. molitor* beetle (Exp. 1, n = 20) or a *P. scaber* woodlouse (Exp. 2, n = 20). Similarly, in experiments 3 and 4 (n = 20 each), the spider was offered as prey a beetle treated either with *P. scaber* defensive secretion (1 equivalent) and water (50 µL) (Exp. 3) or with water only (50 µL) (Exp. 4). After 24 h, all replicates were terminated and predation events were recorded. A dead prey covered in spider silk was deemed preyed upon, whereas a live prey without silk cover was deemed not preyed upon by the spider.

Prediction 2 – spiders reject beetles rendered chemically-defended by treatment with synthetic *P. scaber* secretion metabolites – was tested in experiment 5. With evidence that the synthetic blend (40 ng total) of four *P. scaber* secretion metabolites deterred predation by spiders (see Results), follow-up experiments 6–9 (n = 20 each) tested the effect of each metabolite singly. For each experimental replicate, a synthetic metabolite (40 ng) dissolved in water (50 µL) was applied topically onto a beetle. In all experiments, each spider, beetle, and *P. scaber* was tested only once, with the observer blind to the test stimulus.

### (e) Statistical analysis

All data were analysed in R using Fisher’s Exact Test. Specifically, we compared differential predation of spiders on (*i*) chemically-undefended *T. molitor* beetles (Exp. 1) and chemically-defended *P. scaber* woodlice (Exp. 2), (*ii*) *T. molitor* beetles topically treated with *P. scaber* secretion and water (Exp. 3) or with the four synthetic metabolites **1, 2, 9** and **10** (Exp. 5), and (*iii*) *T. molitor* beetles topically treated with water only (Exp. 4) or with metabolite **1** (Exp. 6), **2** (Exp. 7), **9** (Exp. 8), or **10** (Exp. 9).

## 3. Results

### (a) Identification of defensive metabolites in tegumental gland secretions

The total ion chromatogram [TIC; gas chromatography-mass spectrometry (GC-MS)] of *P. scaber* tegumental gland secretions revealed two distinct metabolites (**1** and **2** in Fig. 1a). The mass spectrum of unknown **1** (Fig. 1b), with an uneven molecular ion (*m*/*z* 203), was indicative of a nitrogen-containing compound, likely with aromatic ring structures. This inference was inspired by *m*/*z* 51, 63, and 76 which are reminiscent of aromatic heterocyclic quinoline. As the molecular weight of **1** (203 Da) was higher than that of quinoline (129 Da), we predicted that **1** has one or more additional functional groups. The presence of a carboxylic acid or a methyl ester functionality was supported by *m*/*z* 143 (203–60), and the presence of a methoxy group by *m*/*z* 171 (203–32). The tailing peak shape of **1** in the TIC (Fig. 1a) further suggested the presence of a polar hydroxyl group. All data combined led us to hypothesize that **1** is a methyl hydroxyquinoline-carboxylate. To test our hypothesis, we purchased 4-hydroxy-quinoline-6-carboxylic acid (**3**), 5-hydroxy-quinoline-3-carboxylic acid (**4**) (both Toronto Research Chemicals, North York, CA), and 8-hydroxy-quinoline-2-carboxylic acid (**5**) (Combi-Blocks Inc., San Diego, USA), and selectively methylated the acid group of all three quinolines using catalytic sulfuric acid in methanol. The methylation product of **5** – methyl 8-hydroxy-quinolene-2-carboxylate – had retention and mass spectral characteristics identical to unknown **1**. Unknown **2** (Fig. 1a), with molecular weight 233 Da (Fig. 1c), was thought of be a homologue of **1** with an additional methoxy functionality. Drawing on the known biosynthesis of **5** via xanthurenic acid (4,8-dihydroxyquinoline-2-carboxylic acid (**6**)) in the kynurenine pathway [30], we proposed two structural candidates for unknown **2**: methyl 8-hydroxy-4-methoxy-quinoline-2-carboxylate (**7**) and methyl 4-hydroxy-8-methoxy-quinoline-2-carboxylate (**8**). Purchased **8** (Combi-Blocks) did not match unknown **2**, but could be used to produce methyl 8-hydroxy-4-methoxy-quinoline-2-carboxylate (Fig. 1c) (see supplemental information), which had retention and mass spectral characteristics entirely consistent with unknown **2**.

**Figure 1.**
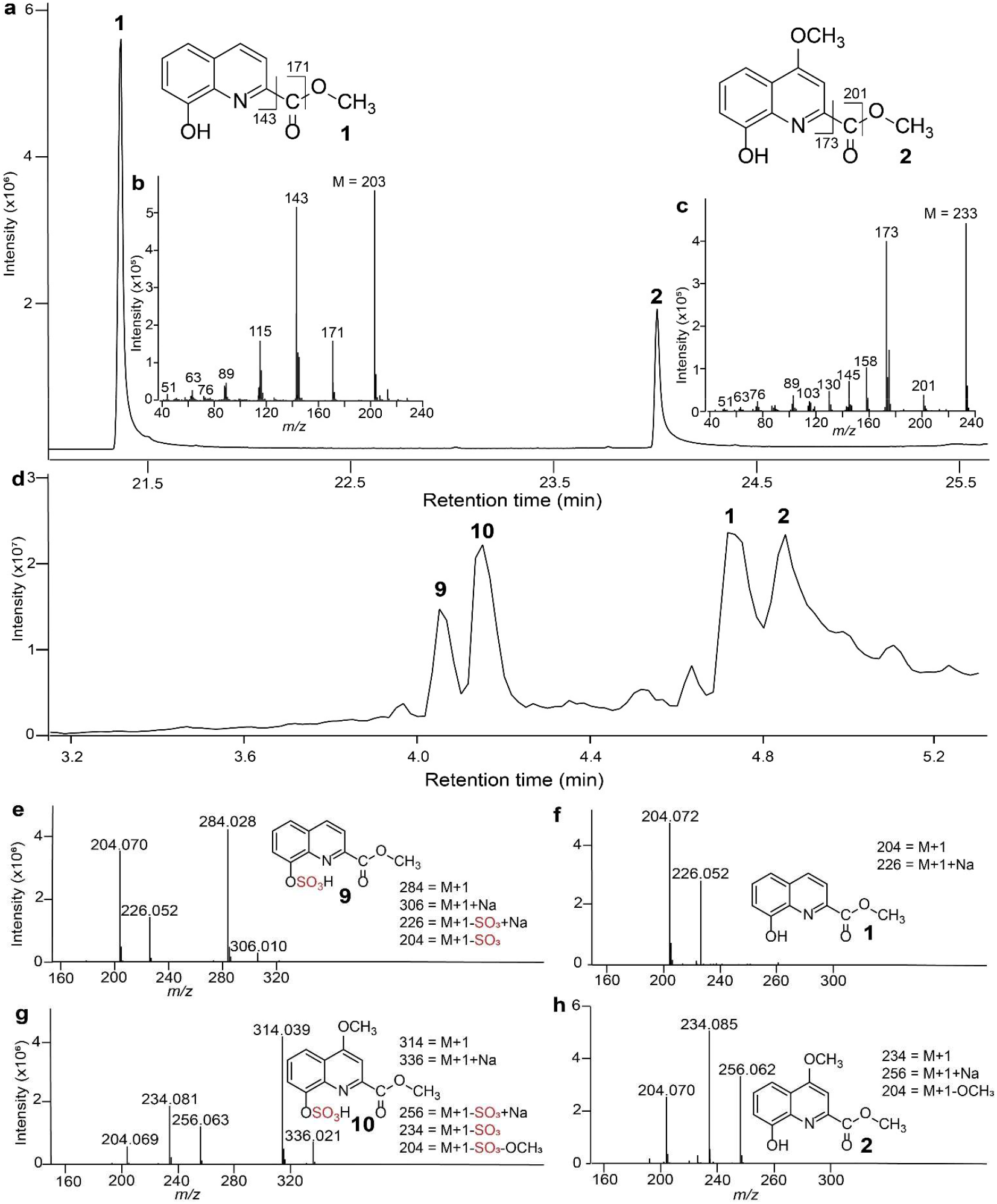
Chromatographic-mass spectrometric analyses of defensive secretions of *Porcellio scaber*. (**a**) Total ion chromatogram (TIC) of gas chromatography-mass spectrometry (GC-MS). (**b, c**) Electron ionization spectra of (**b**) methyl 8-hydroxy-quinoline-2-carboxylate (**1**) and (**c**) methyl 8-hydroxy-4-methoxy-quinoline-2-carboxylate (**2**). (**d**) TIC of high performance liquid chromatography-mass spectrometry (HPLC-MS). (**e-h**) Electrospray ionization spectra of (**e, f**) methyl 8-(sulfooxy)quinoline-2-carboxylate (**9**) and **1**, and of (**g, h**) methyl 4-methoxy-8-(sulfooxy)quinoline-2-carboxylate (**10**) and **2**.

Liquid chromatographic-tandem mass spectrometric (LC/MS-MS) analysis of *P. scaber* secretion extract confirmed the presence of **1** and **2**, and revealed unknown **9** and **10** (Fig. 1d). The retention time differential (0.10 min) of unknown **9** and **10** resembled that of **1** and **2** (0.13 min) suggesting structural similarities. This inference was supported by similar positive ESI spectra of **9** (Fig. 1e) and **1** (Fig. 1f), revealing near identical fragment ions [**9**: *m*/*z* 204.070 (M+1); m/z 226.052 (M+1+Na); **1**: *m*/*z* 204.072; *m*/*z* 226.052]. The additional fragment ions in the spectrum of **9** (*m*/*z* 284.028 (M+1), *m*/*z* 306.010 (M+1+Na)], and the mass differential (79.958 Da) between the molecular ion (+1) of **9** (*m*/*z* 284.028) and **1** (*m*/*z* 204.072), suggested the presence of a -SO_3_ group (79.957 Da) in **9**. Similarly, the ESI spectra of **10** (Fig. 1f) and **2** (Fig. 1h), revealing near identical fragment ions [**10**: *m*/*z* 204.069; *m*/*z* 234.081; *m*/*z* 256.063; *m*/*z* 314.039 (M+1); **2**: *m*/*z* 204.070; *m*/*z* 226.052; *m*/*z* 234.085; *m*/*z* 256.062] suggested structural similarities between **10** and **2**. The additional fragment ions in the spectrum of **10** [*m*/*z* 256.063 (M+1+Na–SO_3_); *m*/*z* 314.039 (M+1); *m*/*z* 336.21 (M+1+Na)], and the mass differential (79.954 Da) between the molecular ion of **10** (*m*/*z* 314.039) and **1** (*m*/z 234.0852), suggested again the presence of an -SO_3_ group in **10**. Drawing on the biosynthetic kynurenine pathway [30], we predicted the sulfooxy group to be at C8, resembling 3-hydroxykynurenine-*O*-sulfate produced by bed bugs [31]. Thus, we synthesized both methyl 8-(sulfooxy)quinoline-2-carboxylate and methyl 4-methoxy-8-(sulfooxy)quinoline-2-carboxylate (see suppl. information), which had retention and mass spectrometric characteristics entirely consistent with unknown **9** and **10**, respectively. With authentic standards of all metabolites in *P. scaber* gland secretion extract at hand, we determined that the tegumental gland secretions of a single *P. scaber* woodlouse contained, on average (1 equivalent), 12 ng of **1**, 18 ng of **2**, 4 ng of **9**, and 6 ng of **10**.

### (b) Behavioural results

Prediction 1 – spiders prey more readily on chemically-undefended *T. molitor* beetles than on chemically-defended *P. scaber* – was supported by data in Exps. 1–4. Spiders preyed on 15 out of 20 beetles (Exp. 1) but preyed on only 6 out of 20 *P. scaber* (Exp. 2) (p = 0.010, Fisher’s Exact Test, Fig. 2). Similarly, spiders preyed on only 6 out of 20 beetles treated with *P. scaber* secretions extracted in water (Exp. 3) but preyed on 16 out of 20 beetles treated only with water (Exp. 4) (p = 0.004, Fisher’s Exact Test, Fig. 2).

**Figure 2.**
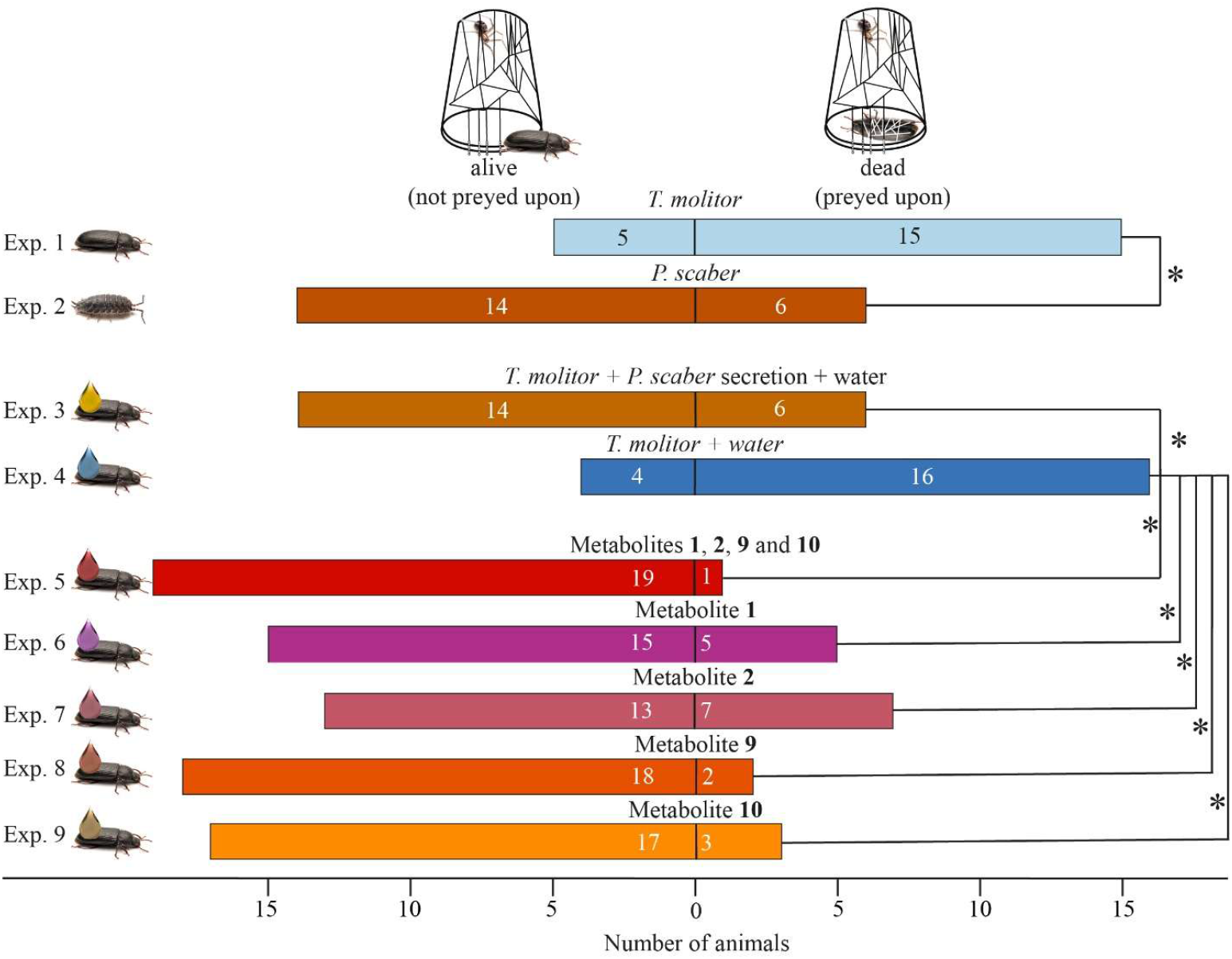
Effects of (*i*) prey type (chemically-defended *Porcellio scaber* woodlice or chemically-undefended *Tenebrio molitor* beetles) and (*ii*) prey treatment (topical application of beetles with *P. scaber* gland secretion, synthetic gland metabolites (40 ng), or a water control) on predation by *Steatoda grossa* false widow spiders. Bars indicate the number of prey being preyed upon and killed (right of 0), or not (left of 0), by the spiders. Metabolites: **1** = methyl 8-hydroxy-quinoline-2-carboxylate; **2** = methyl 8-hydroxy-4-methoxy-quinoline-2-carboxylate; **9** = methyl 8-(sulfooxy)quinoline-2-carboxylate; **10** = methyl 4-methoxy-8-(sulfooxy)quinoline-2-carboxylate. An asterisk (*) denotes a statistically significant difference (Fisher’s Exact Tests; p < 0.05).

Prediction 2 – spiders reject beetles rendered chemically-defended by treatment with synthetic *P. scaber* secretion metabolites – was supported by data in Exp. 5. Spiders preyed on only 1 out of 19 beetles treated with a synthetic blend of **1, 2, 9** and **10** (at 1 woodlouse equivalent) but preyed on 16 out of 20 beetles treated only with water (Exp. 4) (p < 0.001, Fisher’s Exact Test, Fig. 2). Compared to spider predation on water-only-treated beetles (Exp. 4), each synthetic woodlouse metabolite applied onto beetles (at 40 ng) protected beetles from predation (**1** in Exp. 6: 15 (alive) *vs*. 5 (dead), p = 0.001; **2** in Exp. 7: 13 *vs*.7, p = 0.009; **9** in Exp. 8: 18 *vs*. 2, p < 0.001; **10** in Exp. 9: 17 *vs*. 3, p < 0.001, Fisher’s tests, Fig. 2).

## 4. Discussion

Our data support the hypothesis that the evolutionary transition of crustaceans from aquatic to terrestrial habitats prompted the evolution of defensive metabolites against terrestrial predators. We show that the quinoline derivates **1, 2, 9** and **10** in the tegumental secretions of *P. scaber* deter predation by terrestrial spiders. With these chemical and behavioural data, there is now unequivocal evidence that crustaceans, like insects [5,7], are chemically defended against predators [17]. As each of the four quinoline derivates on its own deterred predation by spiders, the blend of all four derivates appears to be a surefire mechanism that ensures protection against a diverse guild of terrestrial predators.

This presentation of multiple behaviour-modifying metabolites is favoured in predator-prey arms races [32] because predators co-evolving with prey may adapt and overcome prey defences [33]. The predator-prey arms race invokes the evolution of novel defensive traits, resulting in the diversification of chemical armaments. Interestingly, chemical defences are thought to diversify less rapidly than the offensive venoms of predators. Venomous predators exert strong resistance selection in their prey populations by removing vulnerable prey, with venom resistance, in turn, amplifying counter-selection for novel offensive agents [34]. On the other hand, evolving deterrence of predators does not require prey death but only a bite predators will remember [32,35]. Predators are often generalists and could switch to other prey while maintaining some offensive potency despite evolving prey defences [32,35]. As well, costly offensive or defensive metabolites that do not accrue substantial fitness benefits are expected to be purged over evolutionary time [34]. Considering both the functional need for varied offensive and defensive metabolites and their production costs, one can expect greater diversity in offensive than in defensive metabolites [32].

Predators with particularly effective co-adaptations to circumvent protective traits of their prey are favoured to specialize on that prey. Such specialized predators of terrestrial isopods are woodlouse spiders in the genus *Dysdera* [36]. *Dysdera* spiders have long chelicera mouthparts that can pierce the hard armour of isopods [37] and help minimize contact with the isopods’ tegumental gland secretions. The heavy and polar metabolites **1, 2, 9** and **10** embedded in the proteinaceous gland secretions of *P. scaber* are likely sensed by tip-pore sensilla at the mouth opening of spiders [38], providing gustatory information on the prey prior to ingestion [39]. Likewise, ants perceive prey chemicals based on receptors on their antennae [21,40]. By secreting distasteful metabolites from their tegumental glands, isopods inform predators about their distastefulness and thereby deter predation. Some predators, however, co-adapted traits that circumvent these prey defences [36]. Studying the chemical defences of diverse terrestrial isopods against their specialist *Dysdera* predators would provide insight into evolutionary co-adaptations in predator-prey arms races [17,36], and would help us understand the selective pressures that have shaped the diversity of defensive mechanisms in isopods. This knowledge would also shed light on the co-evolutionary dynamics – and their effects on biodiversity – between predator and prey that were at play when crustaceans colonized land [41].

With reports that tegumental gland secretions of terrestrial Oniscidea isopods, including those of *P. scaber*, deter predators [19,20, this study], it would be intriguing to investigate the chemical defences of aquatic isopods against aquatic predators, and to compare their metabolites to those identified in this study. To date, it is only known that juveniles of the benthic isopod *Glyptonotus antarcticus* are chemically defended because extracts of juveniles were unpalatable to their sea star predator *Odontaster validus* [42].

There is limited knowledge about the metabolites we identified in *P. scaber* secretions. Compound **2** was not previously known nor were the sulfooxy quinolines **9** and **10**. Reports that compound **1** protects the dytiscid water beetle *Ilybius fenestratus* from frog predation [43] provide evidence for convergent evolution because water beetles secondarily transitioned from terrestrial to aquatic habitats, whereas isopods transitioned from water to land. The corresponding acid of **1**, 8-hydroxy-4-methoxy-quinoline-2-carboxylic acid (**5**), is produced by larvae of the noctuid moth *Spodoptera littoralis* and is believed to serve as an iron-chelating agent, whereas a potential defensive role was not explicitly tested [30,44]. This acid **5** – previously termed ‘quinolobactin’ – functions as a siderophore for *Pseudomonas* bacteria facilitating iron acquisition [45]. Larvae of *S. littoralis* biosynthesize **5** from tryptophan via kynurenine and 3-hydroxykynurenine [30]. As the amount of tryptophan in the diet of these moth larvae determined the amount of **5** they produced [30], it is conceivable that the defensive secretions of *P. scaber* require sufficient tryptophan uptake. In carnivorous cone snails, dietary intake is positively correlated with their venom complexity [46]. Whether *P. scaber* also utilizes dietary precursors for the biosynthesis of its defensive quinoline metabolites remains to be investigated. Locating the biosynthesis site of these quinolines could shed light on the evolution of this defensive trait.

## 5. Conclusion

We have identified four deterrent metabolites (three previously unknown) in the defensive secretion of *P. scaber* supporting the hypothesis that the evolutionary transition of crustaceans from aquatic to terrestrial habitats prompted the evolution of defensive metabolites against terrestrial predators. Our data further demonstrate that some terrestrial crustaceans, like many insects, are chemically defended against predators.

## Ethics

This work did not require ethical approval from a human subject or animal welfare committee.

## Data accessibility

All data and statistical code are provided in the manuscript and in supplementary material which is available online.

## Declaration of AI use

We have not used AI-assisted technologies in creating this article.

## Authors’ contributions

Conceptualization: A.F.; Data curation: A.F.; Formal analysis: A.F., R.G.; Funding acquisition: G.G.; Investigation: A.F., R.G., C.A.R.T., A.D.; Methodology: A.F.; Project administration: A.F.; Resources: G.G., R.G., A.F.; Software: A.F.; Supervision: A.F.; Validation: R.G., A.F.; Visualization: A.F.; Writing – original draft: A.F.; Writing – review & editing: G.G., A.F., R.G. All authors gave final approval for publication and agreed to be held accountable for the work performed therein.

## Acknowledgements

We thank Robert Britton for advice on the synthesis of the sulfooxy quinolines, Hongwen Chen for technical assistance and valuable discussion pertaining to chemical analyses, and Carsten H.G. Müller for discussions on the manuscript. A.F. was supported by an Alexander Graham Bell Scholarship from the Natural Sciences and Engineering Research Council of Canada (NSERC), Graduate Fellowships from SFU, and by the Dr. H.R. McCarthy Bursary.

