## supplementary materials for "Glandular quinoline-derivates protect crustacean woodlice from spider predation"

#### General synthetic procedures

All reactions were performed in oven-dried vessels at ambient temperature and atmosphere unless otherwise specified. Column chromatography was carried out with 230-400 mesh silica gel (E. Merck, Silica Gel 60). Solvents were concentrated and removed via a rotary evaporation. Nuclear magnetic resonance (NMR) spectra were recorded using $CDCl_3$ or $D_2O$ as solvent. Signal positions (δ) are given in ppm from tetramethylsilane (δ 0), and were measured relative to the signal of the solvent ($^1$HNMR: $CDCl_3$: δ 7.26; $D_2O$: δ 4.79; $^{13}$C NMR: $CDCl_3$: δ 77.16). Coupling constants (*J* values) are given in Hertz (Hz). $^1$H NMR spectral data are tabulated in the following order: multiplicity (s, singlet; d, doublet; m, multiplet), coupling constants, number of protons. NMR spectra were recorded on a Bruker Avance 600 equipped with a QNP cryoprobe (600 MHz), a Bruker 500 (500 MHz) or a Bruker 400 (400 MHz). All experiments were run at 298 K. High performance liquid chromatography (HPLC) analyses used an Agilent 1100 high performance liquid chromatograph equipped with a variable wavelength UV-Vis detector. High resolution mass spectra were obtained by electron ionization in positive ion mode on an Agilent 6210 TOF LC/MS, or by electrospray ionization in positive ion mode on a Brucker Maxis Impact. Analytical thin layer chromatography was performed using silica plates, and compounds were visualized at 254 and/or 365 nm ultraviolet irradiation followed by staining with either alkaline permanganate or ethanolic vanillin solution. Commercially available reagents were used throughout without purification unless otherwise stated.

### Specific syntheses

#### Synthesis of methyl 8-hydroxy-4-methoxyquinoline-2-carboxylate (2)

Methyl 4-hydroxy-8-methoxyquinoline-2-carboxylate (0.4 g, 1.715 mmol) was refluxed with potassium iodide (1.139 g, 6.866 mmol) and phosphoric acid (1.681g, 17.10 mmol) to give xanthurenic acid. Xanthurenic acid was then refluxed 24 h in methanol (13 mL) and sulfuric acid (335 µL) at room temperature to give methyl 8-hydroxy-4-methoxyquinoline-2-carboxylate (**2**) [1].

**Syntheses of methyl 8-(sulfooxy)quinoline-2-carboxylate (9) and methyl 4-methoxy-8-(sulfooxy)quinoline-2-carboxylate (10)**

The syntheses of **9** and **10** proceeded in two steps (see below). The first step produced ((fluorosulfonyl)oxy)quinoline precursors which were then treated with cesium carbonate and ethylene glycol to yield **9** and **10**.

**Step 1: Syntheses of methyl 8-((fluorosulfonyl)oxy)quinoline-2-carboxylate and methyl 8-((fluorosulfonyl)oxy)-4-methoxyquinoline-2-carboxylate**

A mixture of tetrahydrofuran (5 ml) and 1,8-diazabicyclo[5.4.0]-undec-7-ene (DBU, see below) was added to an oven-dried vial (10 mL) containing (*i*) methyl 8-hydroxyquinoline-2-carboxylate[2] (100 mg, 0.49 mmol) and [4-(acetylamino)phenyl]imidodisulfuryl difluoride (AISF)[3] [185 mg (0.59 mmol, 1.2 equiv.) or (*ii*) methyl 8-hydroxy-4-methoxyquinoline-2-carboxylate[1] [100 mg (0.42 mmol)] and AISF [161 mg (0.51 mmol, 1.2 equiv.). The volume of DBU in the mixture for (*i*) and (*ii*) was 164 µL (1.0 mmol, 2.2 equiv.) and 143 µL (0.94 mmol, 2.2 equiv.), respectively. After stirring the reaction mixtures 10 min at room temperature, they were diluted with ethyl acetate and washed with 0.5 N HCl and brine. The organic fractions were dried with anhydrous sodium sulfate, filtered and concentrated under reduced pressure. The crude products were purified by silica gel flash chromatography, yielding 125 mg (89%) of methyl 8-((fluorosulfonyl)oxy)-quinoline-2-carboxylate or 110 mg (81%) of methyl 8-((fluorosulfonyl)oxy)-4-methoxyquinoline-2-carboxylate as white solids.

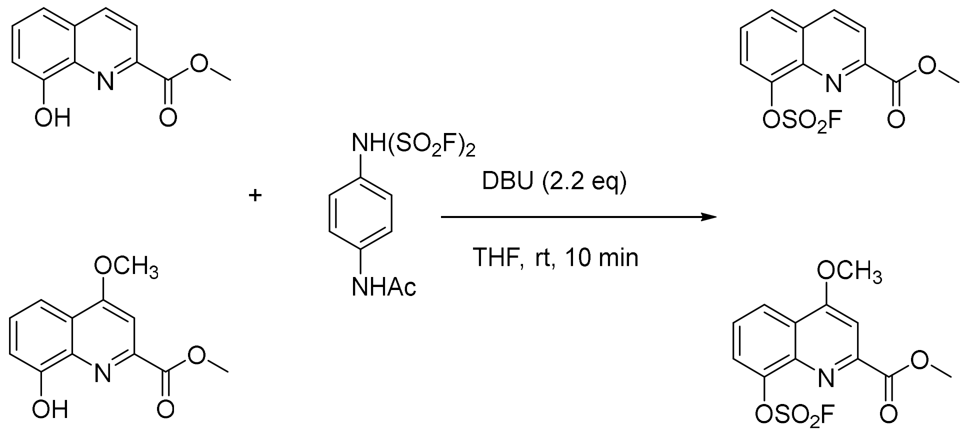


**Methyl 8-((fluorosulfonyl)oxy)-quinoline-2-carboxylate:**

**$^{1}$H NMR:** (400 MHz, $CDCl_3$) δ 8.38 (d, *J* = 8.5 Hz, 1H), 8.29 (d, *J* = 8.6 Hz, 1H), 7.95 (d, *J* = 8.3 Hz, 1H), 7.81 (d, *J* = 7.7 Hz, 1H), 7.71-7.68 (m, 1H), 4.06 (s, 3H)

**$^{13}$C NMR:** (101 MHz, $CDCl_3$) δ 165.4, 149.3, 146.4, 139.8, 137.4, 130.8, 128.5, 128.0, 122.6, 122.2, 53.3

**$^{19}$F NMR:** (376 MHz, $CDCl_3$) δ 41.41 (s, 1F)

**HRMS (ESI)** Calcd. for $C_{11}H_9FNO_5S$: 286.0185 ($[M+H]^+$), found 286.0187

**Methyl 8-((fluorosulfonyl)oxy)-4-methoxyquinoline-2-carboxylate:**

**$^{1}$H NMR:** (400 MHz, $CDCl_3$) δ 8.28 (dd, *J* = 8.4, 1.3 Hz, 1H), 7.78 (d, *J* = 7.7 Hz, 1H), 7.68 (s, 1H), 7.64-7.61 (m, 1H), 4.16 (s, 3H), 4.06 (s, 3H)
**$^{13}$C NMR:** (101 MHz, $CDCl_3$) δ 165.9, 163.5, 150.6, 146.4, 141.0, 126.8, 124.1, 123.0, 122.5, 101.7, 56.6, 53.4
**$^{19}$F NMR:** (376 MHz, $CDCl_3$) δ 41.46 (s, 1F)
**HRMS (ESI)** Calcd. for $C_{12}H_{11}FNO_6S$: 316.0291 ($[M+H]^+$), found 316.0289

**Step 2: Syntheses of methyl 8-(sulfooxy)quinoline-2-carboxylate (9) and methyl 4-methoxy-8-(sulfooxy)quinoline-2-carboxylate (10)**

A mixture of acetonitrile (2 mL) and ethylene glycol [4 µL (0.07 mmol, 1.0 equiv.) or 3.5 µL (0.06 mmol, 1.0 equiv.)] was added to a dry vial (10 mL) containing (*i*) methyl 8-((fluorosulfonyl)oxy) quinoline-2-carboxylate (20 mg, 0.07 mmol) and $Cs_2CO_3$ [46 mg (0.14 mmol, 2 equiv.)] or (*ii*) methyl 8-((fluorosulfonyl)oxy)-4-methoxyquinoline-2-carboxylate (20 mg, 0.06 mmol) and $Cs_2CO_3$ [41 mg (0.12 mmol, 2 equiv.)]. The respective mixtures were stirred 24 h at room temperature.

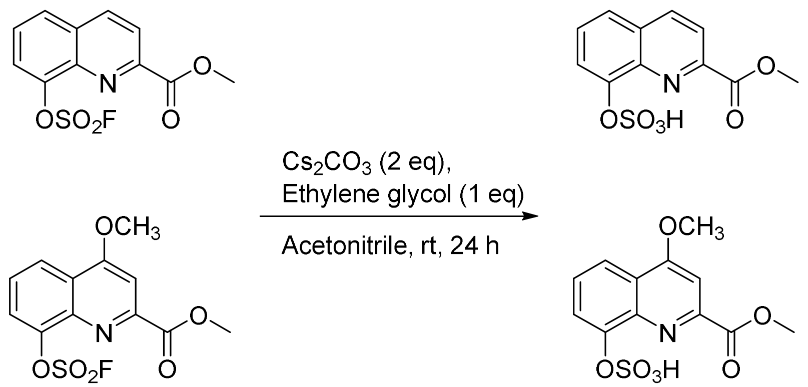


The crude products were purified by HPLC using an Agilent 1100 Series controller connected to a G1361A preparative pump and fitted with a UV detector (set to 254 nm) and a Gemini-NX $C_{18}$ column (50 × 30 mm, 5 µ, 110A). The column was eluted with a 0.4-mL/min flow of a linear solvent (water: acetonitrile) gradient starting from 98:2 and ending with 2:98 after 15 min. Methyl 8-(sulfooxy)quinoline-2-carboxylate)[4] (**9**) (15 mg, 75%) and methyl 4-methoxy-8-(sulfooxy)quinoline-2-carboxylate (**10**) (14 mg, 71%) eluted at retention times 4.56 min and 5.22 min, respectively. Both compounds were white solids.

**Methyl 8-(sulfooxy)quinoline-2-carboxylate:**

**$^{1}$H NMR:** (600 MHz, $D_2O$) δ 8.27 (d, *J* = 8.4 Hz, 1H), 7.95 (d, *J* = 8.5 Hz, 1H), 7.78 (d, *J* = 7.7 Hz, 1H), 7.67 (d, *J* = 8.3 Hz, 1H), 7.59-7.57 (m, 1H), 4.00 (s, 3H)
**$^{13}$C NMR:** (151 MHz, $D_2O$) δ 166.5, 146.9, 146.7, 139.4, 138.4, 130.4, 128.6, 125.1, 121.3, 120.2, 53.2
**HRMS (ESI)** Calcd. for $C_{11}H_{10}NO_6S$: 284.0229 ($[M+H]^+$), found 284.0225

**Methyl 4-methoxy-8-(sulfooxy)quinoline-2-carboxylate:**

**$^1$H NMR:** (600 MHz, $D_2O$) δ 7.70-7.67 (m, 1H), 7.56-7.53 (m, 1H), 7.39-7.35 (m, 1H), 6.97 (s, 1H), 3.94 (s, 3H), 3.80 (s, 3H)
**$^{13}$C NMR:** (151 MHz, $D_2O$) δ 166.2, 162.9, 147.7, 146.6, 139.7, 127.3, 122.6, 119.5, 118.2, 100.6, 56.0, 53.1
**HRMS (ESI)** Calcd. for $C_{11}H_{10}NO_6S$: 314.0334 ($[M+H]^+$), found 314.0328


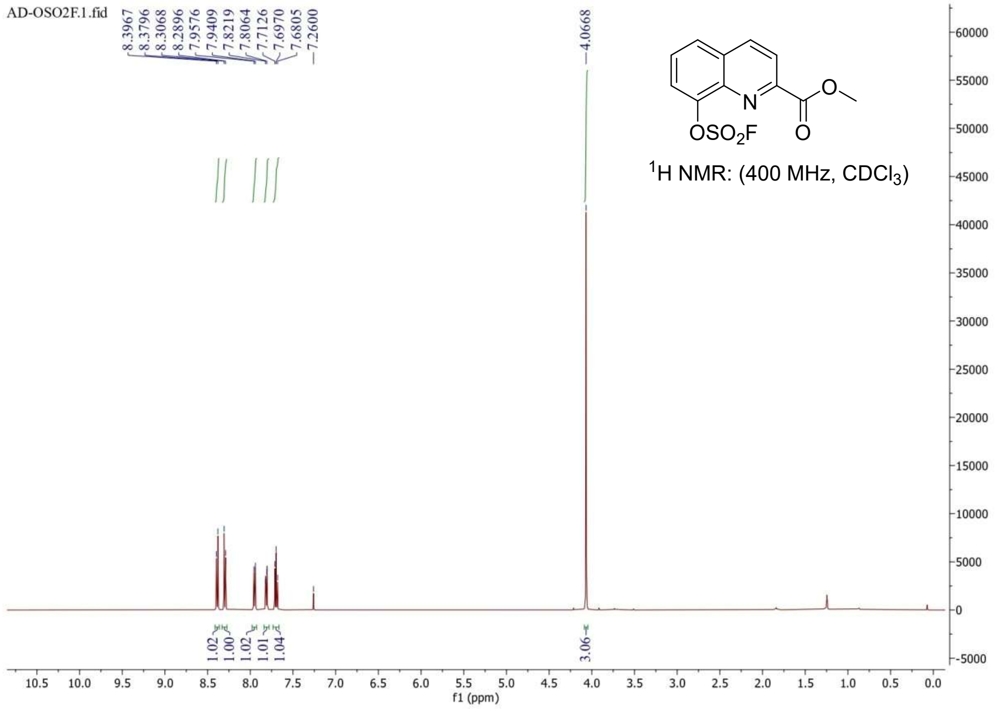

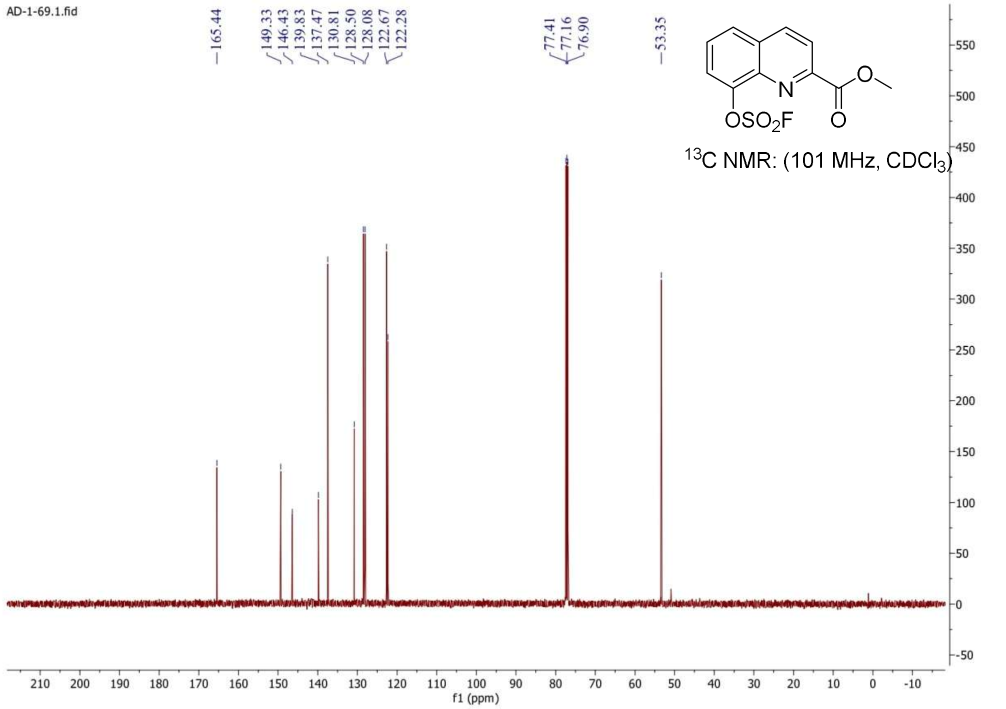
AD-1-69.1.fid
165.44
149.33
146.43
139.83
137.47
130.81
128.50
128.08
122.67
122.28
77.41
77.16
76.90
53.35
N
O
OSO2F
13C NMR: (101 MHz, CDCl3)
f1 (ppm)

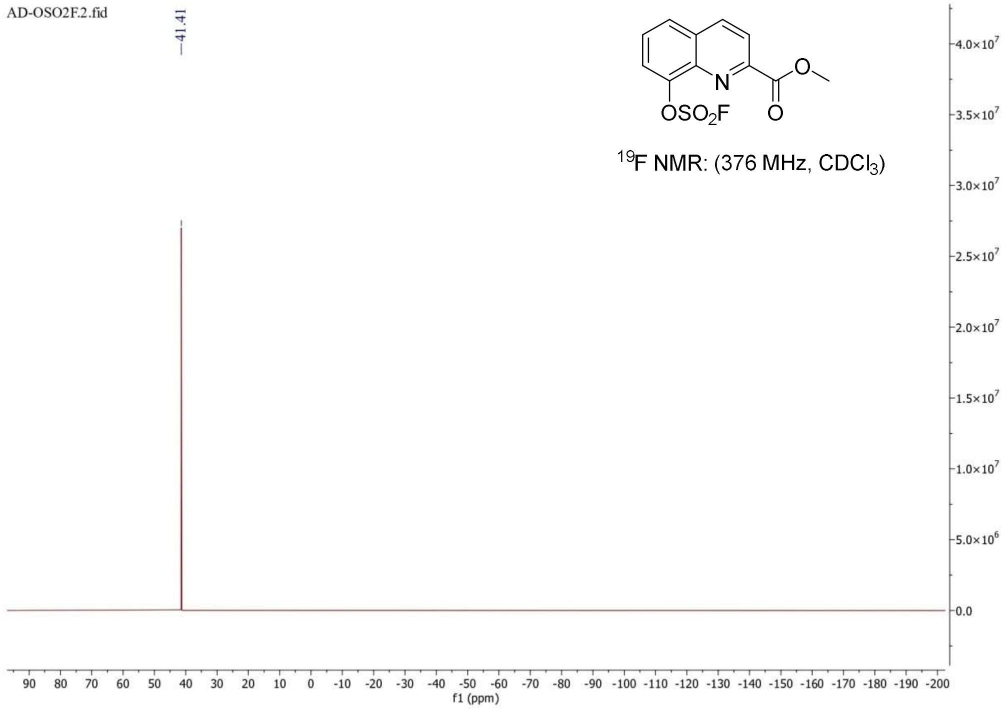
AD-OSO2F.2.fid
41.41
N
O
OSO2F
19F NMR: (376 MHz, CDCl3)
f1 (ppm)

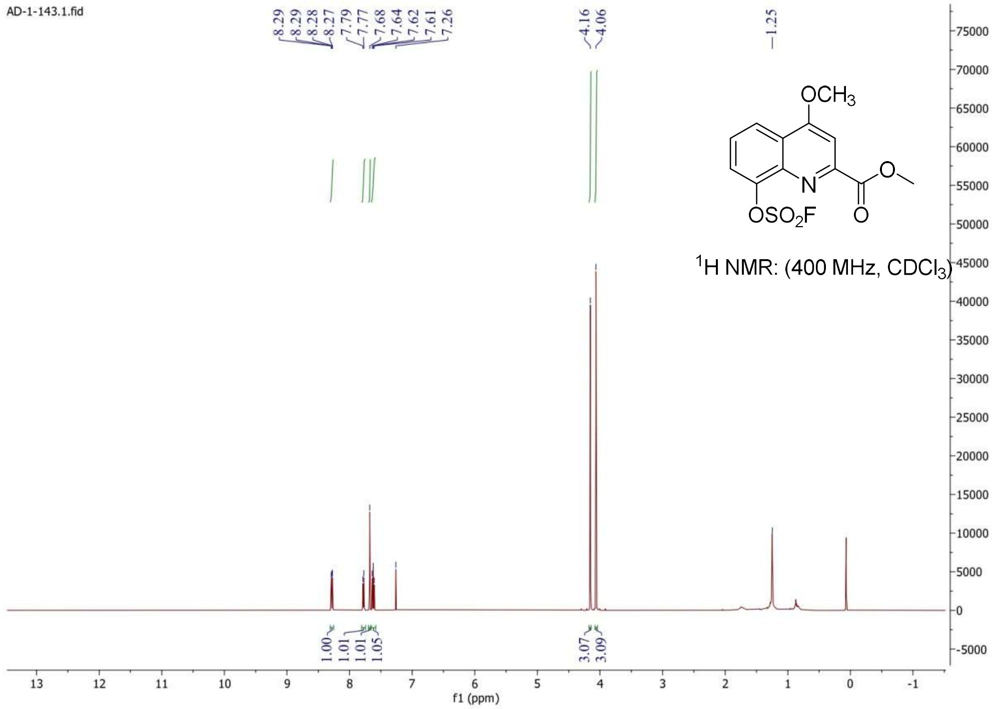
AD-1-143.1.fid
8.29
8.29
8.28
8.27
7.79
7.77
7.68
7.64
7.62
7.61
7.26
4.16
4.06
1.25
OCH3
N
OSO2F
O
O
1H NMR: (400 MHz, CDCl3)
1.00
1.01
1.01
1.05
3.07
3.09
f1 (ppm)

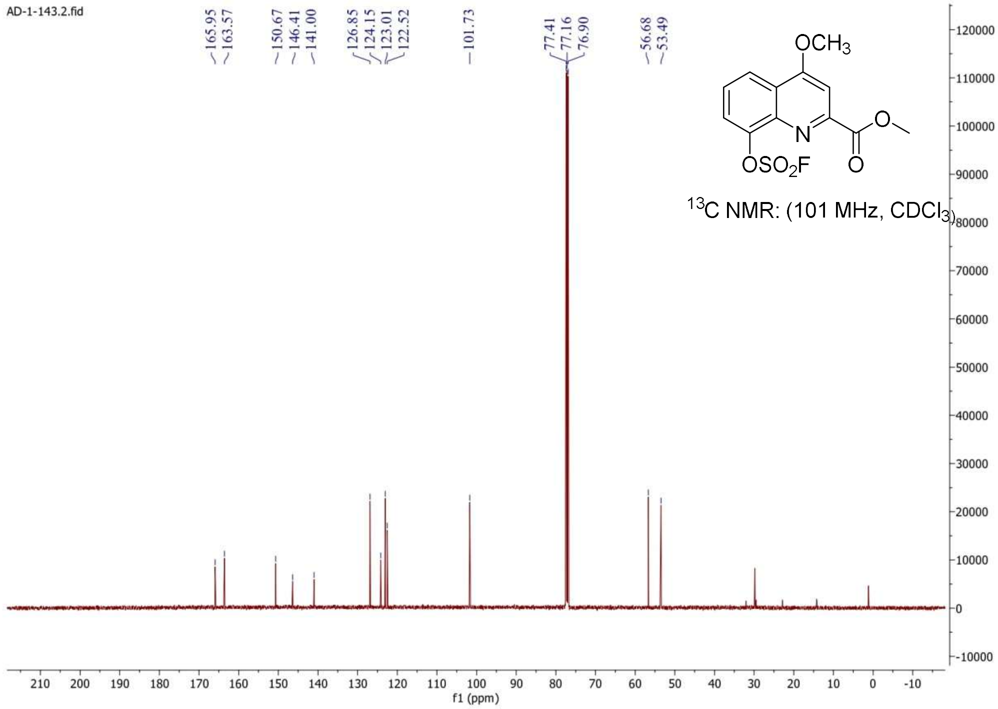
AD-1-143.2.fid
165.95
163.57
150.67
146.41
141.00
126.85
124.15
123.01
122.52
101.73
77.41
77.16
76.90
56.68
53.49
OCH3
N
OSO2F
O
O
13C NMR: (101 MHz, CDCl3)
f1 (ppm)

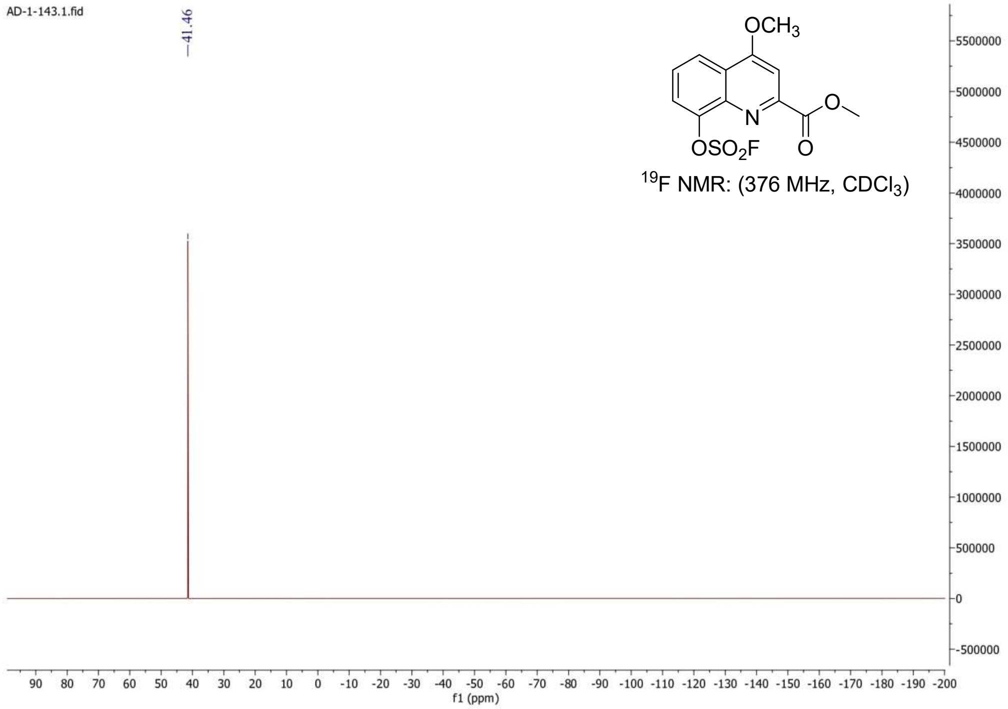

AD-1-143.1.fid
41.46
OCH3
N
OSO2F
O
19F NMR: (376 MHz, CDCl3)
f1 (ppm)


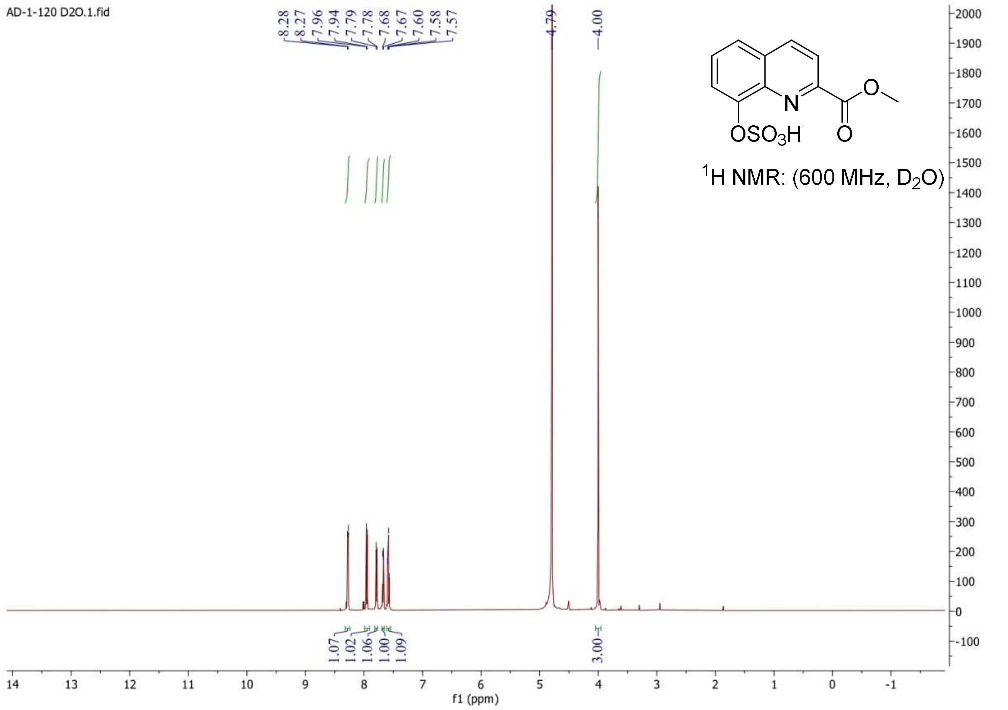

AD-1-120 D2O.1.fid
8.28 8.27 7.96 7.94 7.79 7.78 7.68 7.67 7.60 7.58 7.57
4.79
4.00
N
OSO3H
O
1H NMR: (600 MHz, D2O)
1.07 1.02 1.06 1.00 1.09
3.00
f1 (ppm)

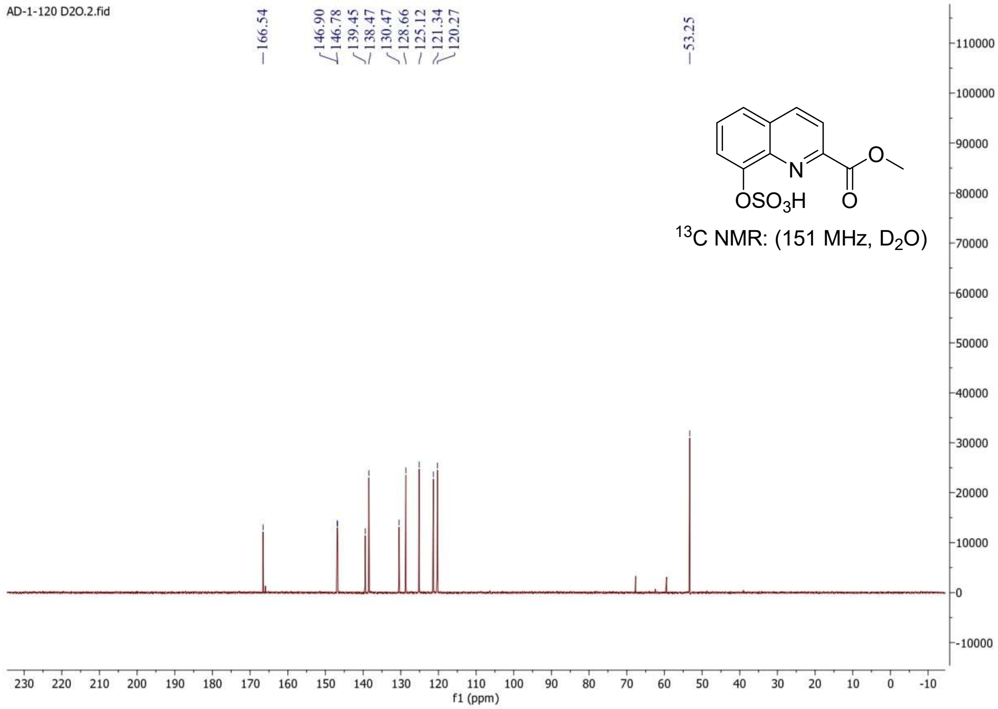
AD-1-120 D2O.2.fid
166.54
146.90
146.78
139.45
138.47
130.47
128.66
125.12
121.34
120.27
53.25
OSO3H
N
O
13C NMR: (151 MHz, D2O)
f1 (ppm)

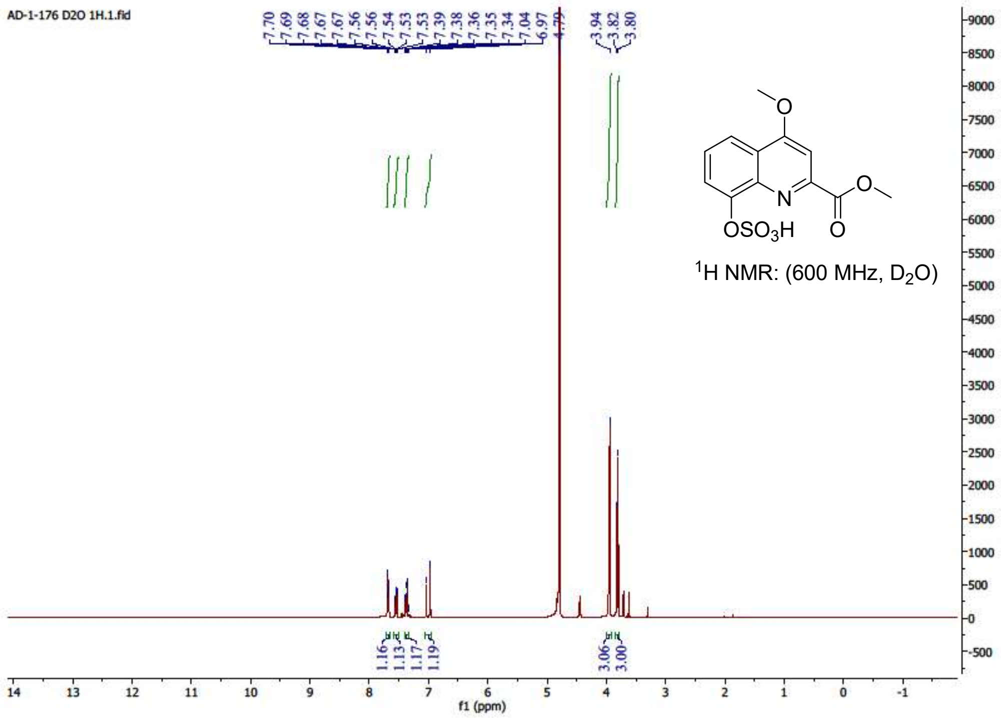
AD-1-176 D2O 1H.1.fid
7.70
7.69
7.68
7.67
7.67
7.56
7.54
7.54
7.53
7.53
7.39
7.38
7.36
7.35
7.34
7.04
6.97
4.79
3.94
3.82
3.80
OSO3H
N
O
1H NMR: (600 MHz, D2O)
1.16
1.13
1.17
1.19
3.06
3.00
f1 (ppm)

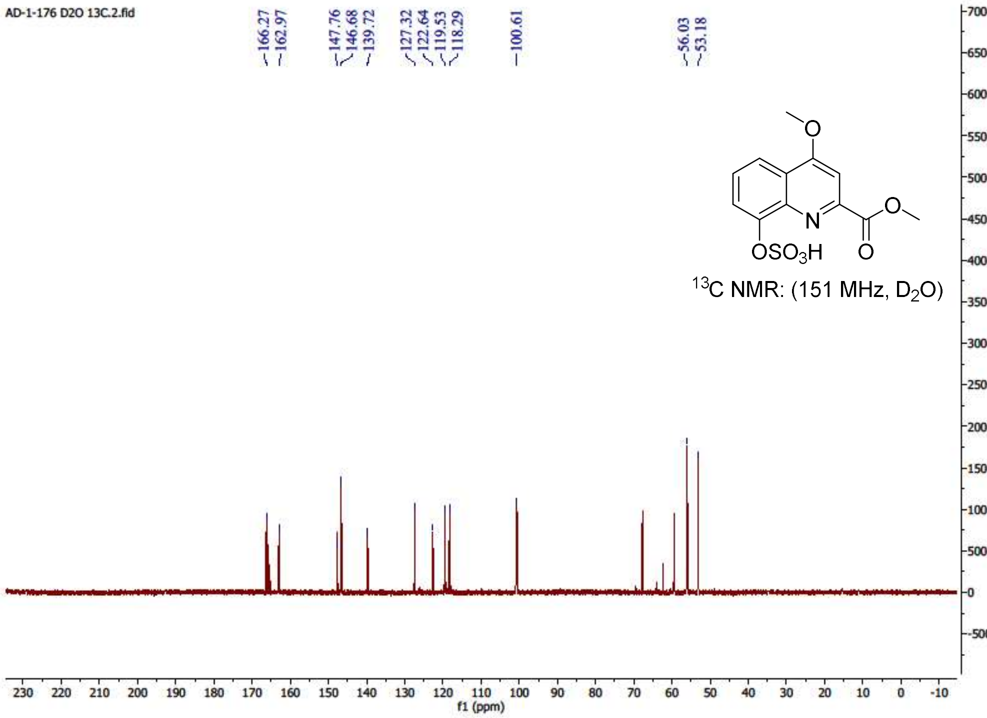
AD-1-176 D2O 13C.2.fid
166.27
162.97
147.76
146.68
139.72
127.32
122.64
119.53
118.29
100.61
56.03
53.18
O
N
O
OSO3H
O
13C NMR: (151 MHz, D2O)
70000
65000
60000
55000
50000
45000
40000
35000
30000
25000
20000
15000
10000
5000
0
-5000
230 220 210 200 190 180 170 160 150 140 130 120 110 100 90 80 70 60 50 40 30 20 10 0 -10
f1 (ppm)
